# CCAN forms the spherical shell-like structure in interphase kinetochores

**DOI:** 10.64898/2026.01.14.699427

**Authors:** Yuji Sakai, Mariko Ariyoshi, Yasuhiro Hirano, Aussie Suzuki, Toru Hirota, Masashi Tachikawa, Tatsuo Fukagawa

## Abstract

The kinetochore is an essential structure for the faithful chromosome segregation during mitosis. The 16-protein complex known as the constitutive centromere-associated network (CCAN) localizes the kinetochore throughout the cell cycle and forms the basis of the kinetochore. Although a structural model of an individual CCAN unit has been proposed through cryo-electron microscopy analysis, little is known about how multiple CCAN units are organized within cells. Here, using *in silico* and *in vivo* analyses, we demonstrate that multiple CCAN units form a spherical, shell-like structure in interphase DT40 kinetochores. CENP-T and CENP-N localize to the CCAN shell, and the CENP-C cupin domain clusters at the center. Molecular dynamics simulations of CCAN assembly suggest that CENP-C contributes to the orientation of CCAN units, leading to the formation of a spherical shell-like structure via CENP-C oligomerization. Super-resolution imaging reveals the spherical shell-like structure at DT40 kinetochores, and knocking out CENP-C abolishes the structure, supporting the reliability of the predicted models. These findings provide important insights into the spatial organization of functional kinetochores within cells.

## Introduction

The kinetochore is a large protein complex formed at the centromere of each chromosome. This ensures accurate segregation of the chromosomes to daughter cells during mitosis and meiosis in all eukaryotes. It is composed of two major complexes: the constitutive centromere-associated network (CCAN) and the KMN (KNL1, Mis12, and Ndc80 complexes) network (*1–6*). In vertebrates, CCAN is represented by a 16-protein complex that forms the basis of interphase kinetochores, which are defined by the histone H3 variant CENP-A (*7–15*). During the late G2 and M phases, CCAN recruits the KMN network, which binds directly to spindle microtubules, forming a functional kinetochore (*2, 3, 16–20*).

Cryo-electron microscopic (EM) analyses have proposed structural models of the human CCAN associated with the CENP-A nucleosome (*6, 21, 22*). While these models provide valuable insights into the molecular organization of an individual CCAN unit, they lack critical structural information for understanding functional CCAN assembly. In particular, the C-terminal region of CENP-C, an essential CCAN protein (*23–28*), was truncated in the proposed CCAN structure models (*21, 22*). Additionally, a significant portion of CENP-C could not be mapped due to its disordered characteristics. Consequently, the contribution of CENP-C to kinetochore organization remains unclear. Furthermore, since each kinetochore contains dozens of CCAN copies, it is largely unknown how multiple CCAN units are organized spatially within cells.

We previously examined the functional roles of CENP-C domains by combining genetics and structural biology using chicken DT40 cells (*29–31*). In these studies, we demonstrated that both the N-terminal KMN binding and CENP-A binding domains of CENP-C are dispensable for mitotic progression. Instead, the CCAN binding domain or the C-terminal cupin domain is essential for CENP-C function. Furthermore, we showed that the vertebrate CENP-C cupin domain exhibits oligomerization activity, which is essential for CENP-C function (*31*). A fusion of the CCAN-binding domain and the C-terminal cupin domain (called “Mini-CENP-C”) rescues the CENP-C knockout phenotype. This suggests that an essential role of CENP-C is organizing CCAN via direct binding and oligomerization (*6, 31*). However, the contribution of CENP-C oligomerization properties to the higher-order assembly of multiple CCAN units and the actual structure of CCAN formed within cells remain unclear.

In this study, we created coarse-grained (CG) models of the Mini-CENP-C and CCAN subcomplexes. We then performed a molecular dynamics (MD) simulation of CCAN assembly to gain insight into the structure of the kinetochore. Combined with mutational analysis using chicken DT40 cells, our MD simulations revealed that CENP-C promotes face-to-face orientation between CCAN units. The simulations also suggest that the CCAN units assemble into a spherical, shell-like structure with the CENP-C cupin domains clustered at the center. Finally, super-resolution imaging revealed that CCAN forms a spherical, shell-like structure in interphase DT40 kinetochores, and that CENP-C is required for formation of the structure. These findings provide important insights into the spatial organization of the functional kinetochore within cells.

## Results

### CENP-C contributes to the organized orientation of CCAN units in the presence of DNA

To investigate the role of CENP-C cupin domain oligomerization in the formation of higher-order kinetochore structures, we performed MD simulations on a simplified system consisting of two CCAN units bound to a CENP-C dimer. We first generated a dimeric chicken CCAN structural model as the initial model for MD simulations by combining AlphaFold3 (AF3) predictions (*32*) with additional model building (Figs. 1A and S1A, see Methods). To reduce the computational cost of MD simulations while retaining essential CENP-C functions, we used Mini-CENP-C, in which CCAN binding domain (CBD, amino acids (aa) 166-324) is fused with a C-terminal cupin-containing region (aa 677-864), instead of full-length CENP-C (Fig. 1A, left). Previous studies have validated that Mini-CENP-C retains essential CENP-C functions (*31*). Mini-CENP-C contains a CENP-L/N binding motif and a cupin domain connected by an intrinsically disordered region (IDR). Thus, two CCAN units are expected to position themselves in close proximity through cupin-mediated dimerization (Fig. 1A, right). We generated AF3-predicted structural models of the CCAN unit containing the N-terminal half of the Mini-CENP-C CBD (amino acids [aa] 166-235), which includes the CENP-L/N binding motif. We also generated a model of the dimeric Mini-CENP-C cupin domain (aa 706-864) (Fig. S1A). The IDR in each Mini-CENP-C fragment was excluded from the prediction due to low confidence (Fig. 1A, right). Additionally, a 40 bp DNA fragment was included in the AF3 prediction of the CCAN unit, because the CCAN complex forms a continuous DNA-binding interface on its inner surface (*21, 22*). The predicted chicken CCAN model closely resembles to the cryo-EM structure of the human CCAN core complex (*21, 22*), supporting the reliability of the predicted structure (Fig. S1A, upper). Finally, the two CCAN units and the dimeric cupin domains were connected via modeled Mini-CENP-C IDRs, resulting in a macromolecular assembly with an overall dimension of 50 nm (Fig. 1A and Fig. S1B). Coarse-grained (CG) MD simulations based on the Martini model are effective for systems of this scale (*33, 34*).

**Figure 1.**
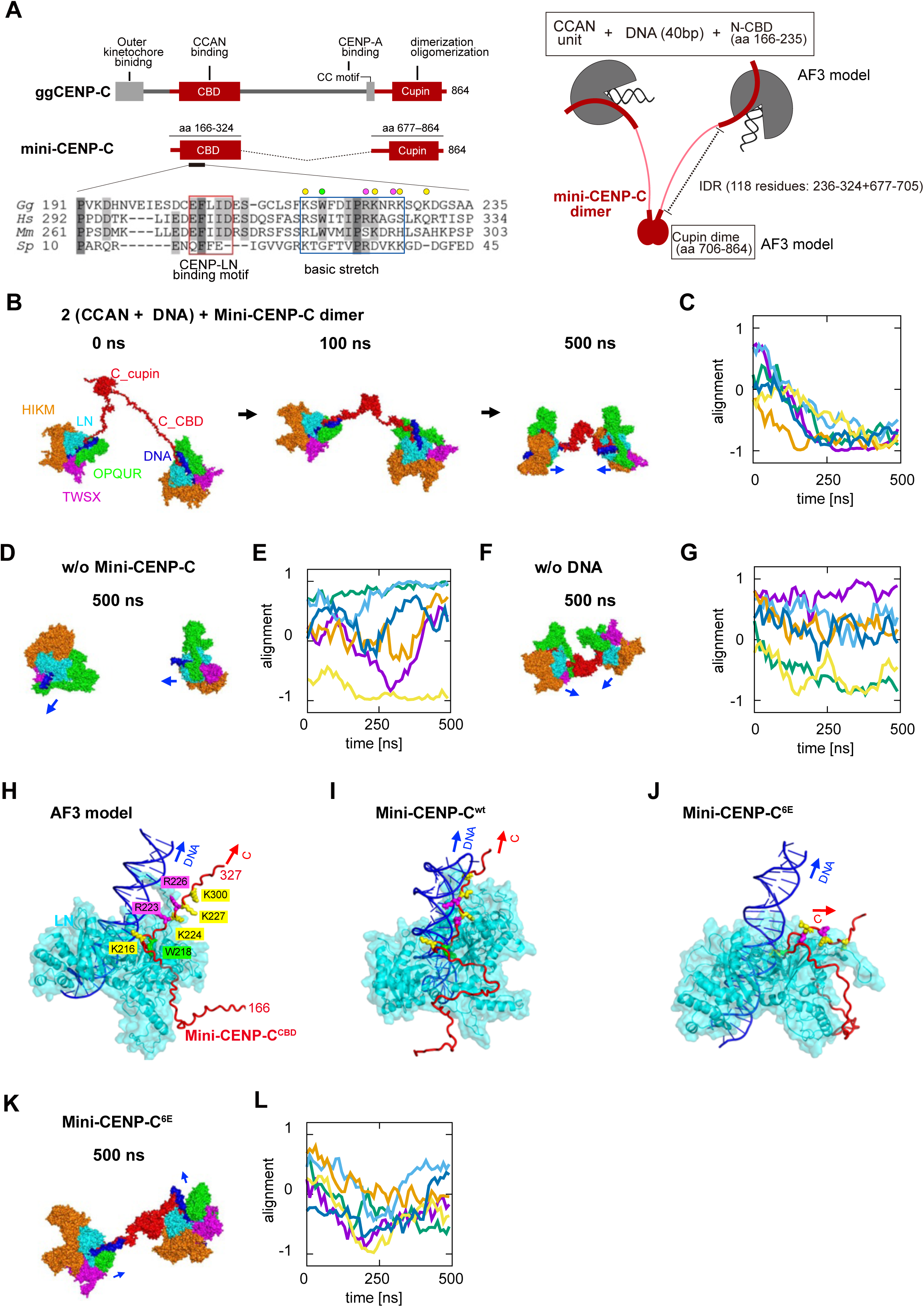
MD simulation of two CCAN units linked by a Mini-CENP-C dimer. (A) Left, schematic representation of the functional regions of chicken CENP-C. Mini-CENP-C comprises of the CCAN-binding domain (CBD; aa 166-324) and the C-terminal cupin domain (aa 677-864). The conserved CENP-L/N binding motif and basic stretch within CENP-C CBD are indicated in the multiple sequence alignment. The residues mutated in the 6E mutant are indicated by colored circles: yellow, K216/224/227/K230; magenta, R223/226; green, W218. Right, schematic of the initial model used for MD simulations, consisting of two CCAN core units, with or without double stranded DNA (40 bp), bound to dimeric Mini-CENP-C. The N-terminal half of CBD (N-CBD) and the globular cupin domain are connected by an intrinsically disordered region (IDR). (B) Representative CG MD snapshots of two CCAN units with DNA linked by a Mini-CENP-C dimer at 0, 100, and 500 ns. CCAN subcomplexes (CENP-H/I/K/M, -L/N, -O/P/Q/U/R, and -T/W/S/X), Mini-CENP-C, and DNA are shown in orange, cyan, green, magenta, red, and blue, respectively. Blue arrows indicate the directional vectors of the DNA axis in the CCAN units. (C) Temporal changes of relative orientation of two CCAN units calculated from six independent simulation using different initial models. Initial structures and representative structures at 500 ns are shown in Supplementary Figures S1C and S1D, respectively. (D) Structures of two DNA-bound CCAN units simulated in the absence of Mini-CENP-C at 500 ns. Color scheme as in panel (B). (E) Temporal changes of relative orientation of two CCAN units calculated from six independent simulations performed without Mini-CENP-C. Representative structures at 500 ns are shown in Supplementary Figure S1E. (F) Structures of two CCAN units lacking DNA but linked by a Mini-CENP-C dimer at 500 ns. Color scheme as in panel (B). (G) Temporal changes of relative orientation of two CCAN units calculated from six independent simulations in the absence of DNA. Representative structures at 500 ns are shown in Supplementary Figure S1F. (H) AlphaFold3-predicted structural model of the CENP-L/N subcomplex bound to 28 bp DNA and CENP-C CBD (aa 166–237). The N-terminal half of Mini-CENP-C CBD (aa 166-237) was included. CENP-L/N is shown as cyan, DNA as blue, and CENP-C as red in panels (H–J). Blue and red arrows indicate directional vectors of DNA helical axis and the main chain path of the basic stretch in Mini-CENP-C (H-J) (I) Structure after 100 ns of all-atom MD simulation using CENP-C initiated from the model shown in panel(H). (J) Structure after 100 ns of all-atom MD simulation using the Mini-CENP-C^6E^ mutant, initiated from the same starting model shown in panel (H). (K) Structures of two CCAN units with DNA linked by the Mini-CENP-C^6E^ mutant at 500 ns. Color scheme as in panel (B). (L) Temporal changes of relative orientation of two CCAN units calculated from six independent simulations with the basic patch mutant Mini-CENP-C. Representative structures at 500 ns are shown in Fig. S1G.

First, using the initial model, we simulated how two CCAN units tethered by Mini-CENP-C settle into equilibrium states (Fig. 1B, Supplementary Movie S1). During the initial phase, the extended IDR in each Mini-CENP-C fragment underwent thermal contraction, shrinking from 20 nm to 7-8 nm and drawing the two CCAN units closer together (100 ns, Fig. 1B). Subsequently, the initially random orientation of the CCAN units rearranged while maintaining their distance, resulting in a face-to-face orientation (500 ns, Fig. 1B). No direct interaction between the CCAN units was observed during the simulation. To investigate whether the equilibrium orientation of the CCAN units depends on the initial configuration, we initiated simulations from multiple CCAN-Mini-CENP-C models with different orientations (Figs. S1B-C). For each simulation, we quantified the temporal changes in CCAN orientation, which is defined by the relative directional vectors between the two DNA fragments, each of which is bound to one CCAN unit (Fig. 1B, 500 ns) (see Methods section). Despite the varied initial states, all simulations converged on a face-to-face orientation of the two directional vectors, (i.e., the CCAN units) in the equilibrium state (Figs. 1C and S1C-D). In contrast, in the absence of Mini-CENP-C, the two CCAN units did not approach each other. Instead, they diffused independently and moved apart without aligning their orientations (Fig. 1D, Supplementary Movie S2). We performed simulations using six different CCAN orientations in the absence of Mini-CENP-C; however, no face-to-face alignment was observed, and the orientation of the CCAN units remained random (Figs. 1E and S1E). These results suggest that Mini-CENP-C drives the orientational alignment of the CCAN units by promoting a face-to-face orientation that would not arise spontaneously.

Next, we examined how DNA influences the face-to-face orientation of the CCAN, which is mediated by Mini-CENP-C. We performed simulations of the CCAN-Mini-CENP-C complex without the DNA fragments and found that the two CCAN units remained in close proximity (Fig. 1F, Supplementary Movie S3). However, in five additional independent simulations that started from different CCAN configurations and excluded DNA, the relative orientations of the two CCAN units were more diverse than those observed in the presence of DNA (Figs. 1G and S1F). Therefore, DNA likely imposes additional constraints that restrict CCAN motion and stabilize its face-to-face orientation mediated by Mini-CENP-C.

We investigated the role of DNA in the organization of the CCAN unit mediated by Mini-CENP-C. In the AF3-predicted structure, the DNA contacts a cluster of positively charged residues (K216/R223/R226) located near the CENP-L/N-binding motif of Mini-CENP-C (Figs. 1A and 1H). This contact was not observed in the cryo-EM structures of the human CCAN complex (*21, 22*). CG MD simulations showed that the CENP-C-DNA binding was stably maintained for at least 500 ns (Fig. 1B and S1D). This CENP-C region contains an extended basic stretch with additional potential DNA-binding residues (Fig. 1A). Therefore, we performed MD simulations using a complex model with mutations introduced to this region (W218A/R223,226E/K216,224,227,230E). Hereafter, these mutations are referred to as 6E. Notably, these CENP-C residues are conserved in other species (Fig. 1A). To analyze CENP-C–DNA interactions more precisely, we performed all-atom MD simulations instead of CG MD simulations. We used a model containing Mini-CENP-C CBD (aa 166–237), CENP-L/N, and DNA. Since the CENP-L/N subcomplex plays a pivotal role in forming a robust DNA-binding grip within the CCAN-DNA complex, CENP-L/N was selected as a representative CCAN subcomplex. In the simulation with the wild-type Mini-CENP-C, DNA binding was stable, and the basic stretch of Mini-CENP-C extended along the DNA axis (Fig. 1I). In contrast, during the simulation with the Mini-CENP-C^6E^ mutant, the basic stretch failed to maintain DNA association and moved away from the DNA. This suggests that the basic stretch of CENP-C stabilizes its interaction with DNA (Fig. 1J).

Finally, we examined the effect of Mini-CENP-C^6E^ on the organization of CCAN units in full CG MD simulations. In simulations containing the CCAN-Mini CENP-C^6E^ complex, the face-to-face orientation of the two CCAN units was disrupted (Fig. 1K, Supplementary Movie S4). We performed additional five independent simulations starting from different CCAN initial models, with Mini-CENP-C^6E^. In all cases, the simulations showed a lower degree of orientational convergence than those using the wild-type Mini-CENP-C, suggesting that the 6E mutations destabilize the orientational ensemble of the CCAN units (Figs. 1L and S1G).

Based on these simulation results, we conclude that dimeric CENP-C imposes spatial constraints on the organization of two CCAN units. Specifically, CENP-C’s DNA binding via the conserved basic stretch is crucial for arranging the two CCAN units in a defined face-to-face orientation.

### CG MD simulations reveal that multiple CCAN units form a spherical shell-like structure with CENP-C clustered at its center

Each kinetochore contains several dozen copies of CCAN (*35, 36*), which may form clusters within cells. We analyzed how multiple CCAN units assemble with CENP-C using CG MD simulations. For these simulations, we generated an initial model consisting of eight CCAN units, each bound to 40 bp of DNA, and four Mini-CENP-C dimers (8(CCAN+DNA) +4Mini-CENP-C dimer, 0 ns) (Fig. 2A, See Methods). In this model, the eight CCAN units were positioned randomly at nearly uniform radial distances (160 ∼ 180Å) from the center formed by the oligomerized cupin domains of Mini-CENP-C. The oligomer model of four CENP-C cupin dimers (eight cupin monomers) was generated based on the oligomer interface observed in the previously reported crystal structure (*31*).

**Figure 2.**
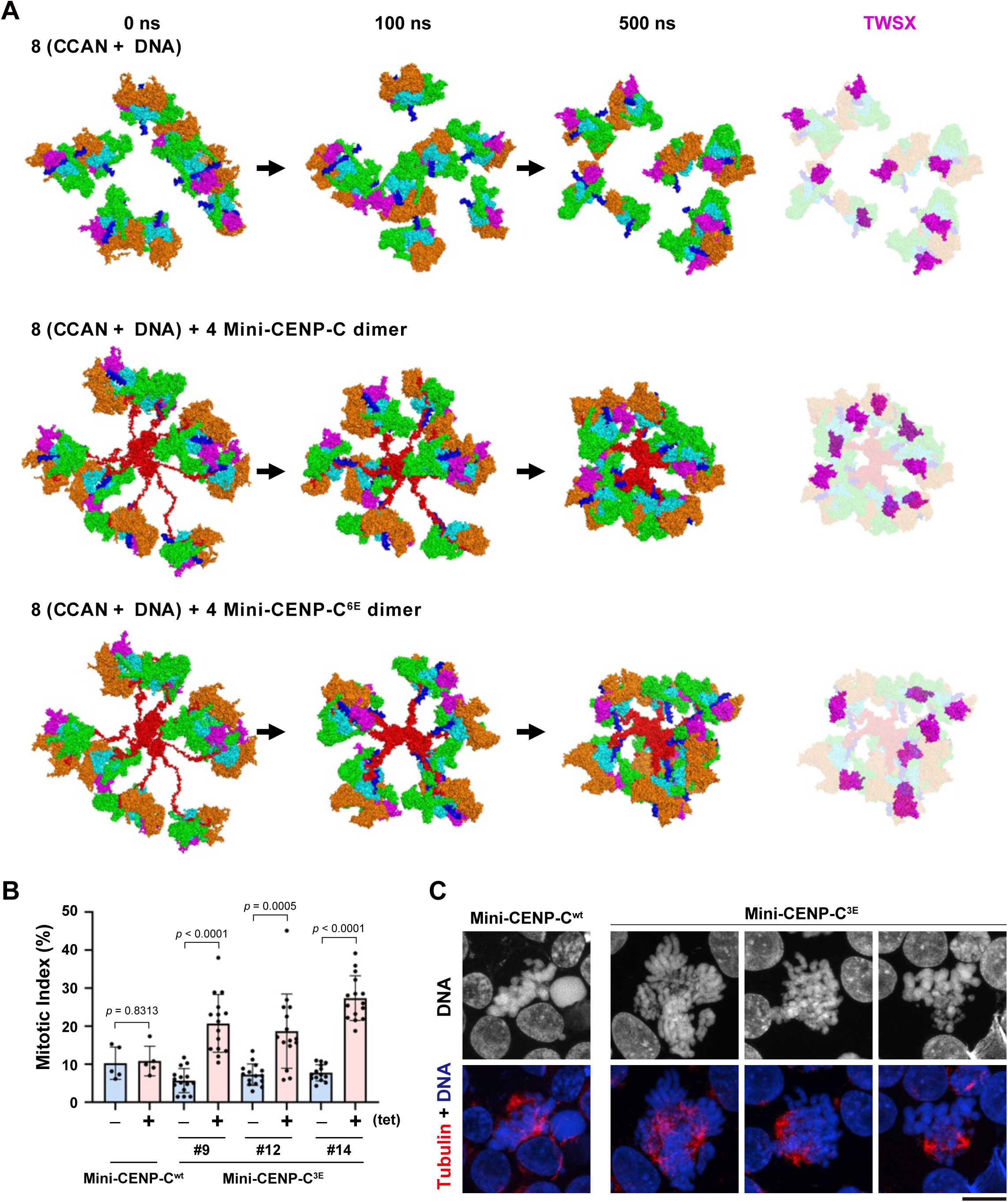
MD simulation of eight CCAN units with DNA linked by four Mini-CENP-C dimers. (A) Top, representative CG MD snapshots of eight CCAN units including DNA, simulated in the absence of Mini-CENP-C at 0, 100, and 500 ns. The right panel highlights the T/W/S/X subcomplexes from the 500 ns structure. Color scheme as in Fig. 1B. Middle, representative CG MD snapshots of eight DNA-bound CCAN units linked by four Mini-CENP-C dimers at 0, 100, and 500 ns. The right panel highlights the T/W/S/X subcomplexes from the 500 ns structure. Color scheme as in Figure 1B. Bottom, representative CG MD snapshots of eight CCAN units with DNA linked by four Mini-CENP-C^6E^ dimers at 0, 100, and 500 ns. The right panel highlights the T/W/S/X subcomplexes from the 500 ns structure. Color scheme as in Figure 1B. (B) Mitotic index of cKO-CENP-C DT40 cells expressing Mini-CENP-C^wt^ or Mini CENP-C^3E^ in the presence (tet+) or absence (tet-) of tetracycline. Three different lines of Mini CENP-C^3E^-expressing cells were used for the quantification. Mitotic cells were judged by spindle morphology. Statistical significance was tested using unpaired two-tailed Welch’s *t*-test. (C) Representative images of mitotic cKO-CENP-C DT40 cells expressing Mini-CENP-C^wt^ or Mini CENP-C^3E^ in the presence of tet. Cells were stained by an anti-α-tubulin (red). DNAs were stained by DAPI (blue). Scale bar, 10µm.

In the absence of Mini-CENP-C, the CCAN units did not assemble into a defined orientation relative to each other, even in the presence of DNA. This is consistent with observations from CG MD simulations using two isolated CCAN units (Figs. 1D and 2A, upper panel, Supplementary Movie S5). However, in a CG MD simulation of an 8(CCAN+DNA)+4Mini-CENP-C dimer model, the eight CCAN units were drawn toward the octameric Mini-CENP-C cupin core. The DNA in each CCAN unit was oriented toward the Mini-CENP-C cupin domain and was arranged in a defined spatial organization (Fig. 2A, middle panel, Supplementary Movie S6). Consistent with the dimeric CCAN-DNA-Mini-CENP-C complex simulation, the IDRs of all eight Mini-CENP-C fragments converged to a similar distance of 7–8 nm (Fig. S2A). In the resulting structure, the CCAN units formed a spherical shell-like organization. Intriguingly, the CENP-T/W/S/X and CENP-L/N subcomplexes were positioned toward the outer region of the spherical structure. Meanwhile, the CENP-C cupin domains remained assembled around the central region, maintaining the oligomeric interface present in the initial model (Fig. 2A middle panel and S2B).

In our MD simulations, we showed that DNA binding of Mini-CENP-C through its basic stretch participates in the face-to-face orientation of two CCAN units tethered by a Mini-CENP-C dimer. Therefore, we examined how the basic stretch contributes to the spherical, shell-like structure of multiple CCAN units. A simulation using Mini-CENP-C^6E^ produced an unorganized assembly of CCAN-DNA units, even though they were drawn toward the cupin core. Consequently, the CENP-T/W/S/X and CENP-L/N subcomplexes adopted a more random pattern and deviated from the spherical outer region compared to simulations using wild-type Mini-CENP-C (Fig. 2A lower panel, Supplementary Movie S7). Taken together, these simulations suggest that multiple CCAN units assemble into a spherical, shell-like organization, in which the units are oriented toward the cupin core, and that DNA binding by Mini-CENP-C plays a key role in establishing this higher-order CCAN assembly.

### CENP-C-DNA interaction is critical for kinetochore function

MD simulations suggest that the CENP-C-DNA interaction is essential for proper CCAN organization. To experimentally test this prediction, we examined the functional importance of the basic stretch of CENP-C in chicken DT40 cells. For this experiment we used a tetracycline (tet)-inducible conditional CENP-C knockout (cKO-CENP-C) cell line (*29, 37*). The cKO-CENP-C cells died after mitotic arrest in the presence of tet; however, expressing Mini-CENP-C^wt^ (wild-type Mini-CENP-C) rescued the phenotype (*31*). Consistent with the previous results, the mitotic index (less than 10%) did not increase in cKO-CENP-C cells expressing Mini-CENP-C^wt^ after the addition of tet (Fig. 2B). However, expression of Mini-CENP-C with K216E/R223E/R226E (3E) mutation in the basic stretch failed to rescue mitotic accumulation in cKO-CENP-C cells after the addition of tet (Fig. 2B). We tested three independent clones of cKO-CENP-C cells expressing Mini-CENP-C^3E^, and the mitotic index increased by 10-20% in all clones after the addition of tet (Fig. 2B).

In addition to the increased mitotic index observed in cKO-CENP-C cells expressing Mini-CENP-C^3E^, abnormal mitotic cells were frequently observed in these clones. Well-aligned chromosomes with bipolar spindles were observed in cKO-CENP-C cells expressing Mini-CENP-C^wt^. However, multi-polar spindles and unaligned chromosomes were observed in cells expressing Mini-CENP-C^3E^ (Fig. 2C). Based on these results, we conclude that the CENP-C-DNA interaction via the basic stretch is critical for CENP-C function.

### The spherical shell-like structure is observed in DT40 kinetochores

Our MD simulations demonstrate that multiple copies of CCAN units form a spherical shell-like structure held together by the oligomerization of the CENP-C cupin domain (Fig. 2A). To investigate the existence of a structure similar to our simulated kinetochore model in DT40 cells, we examined the localization of CENP-T and the C-terminal region of CENP-C. These are predicted to reside in the outer and inner regions of the kinetochore structure, respectively. To this end, we immunostained DT40 cells in which endogenous CENP-T or CENP-C was replaced with mScarlet-HA-SNAP–CENP-T (this study) or CENP-C-GFP (*30*), respectively, with anti-CENP-T or anti-GFP antibodies. Because high-resolution imaging is required to resolve the detailed architecture of the kinetochore, we used 12x expansion microscopy (ExM) for observations (*38*). Using ExM, we observed CENP-T as a spherical, shell-like structure with no signal detected in the central region (Figs 3A and S3, see XZ and YZ plane images). In striking contrast, the C-terminal region of CENP-C appeared as discrete, evenly distributed dot-like structures throughout the entire kinetochore (Figs 3B and S3, see XZ and YZ plane images). The CENP-T spherical shell-like structure was slightly larger than the CENP-C structure, though the difference was not statistically significant (Fig. 3C). This suggests that the CENP-T shell surrounds the CENP-C core, which is consistent with models from our MD simulations.

**Figure 3.**
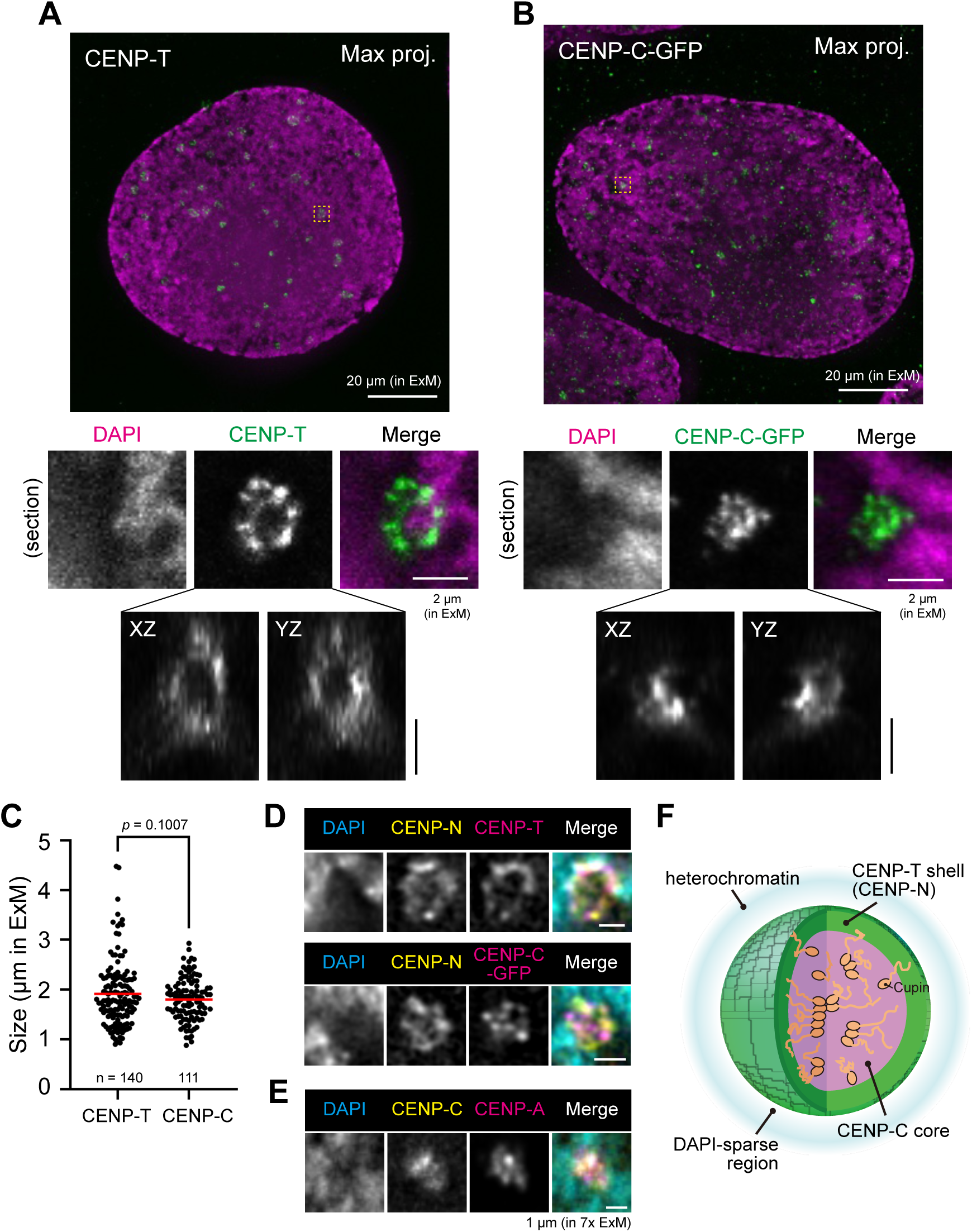
The CCAN forms a shell-like structure with CENP-C as its core. (A) CENP-T forms a shell-like structure. DT40 cells expressing mScarlet-HA-SNAP-CENP-T were immunostained with anti-CENP-T antibody and counterstained with DAPI, followed by expansion microscopy. Representative deconvolved ExM images are shown. The top panel displays a maximum intensity projection of a representative cell, with the yellow boxed region enlarged in the bottom panel. The three-dimensional CENP-T structures are shown in the XZ and YZ planes. Scale bars, 20 µm (left) and 2 µm (right) in ExM. (B) Localization of the C-terminus of CENP-C at the kinetochore core. DT40 cells expressing CENP-C-GFP were immunostained with anti-GFP antibody to visualize the C-terminus of CENP-C. Representative ExM images are shown as in panel A. Scale bars, 20 µm (left) and 2 µm (right) in ExM. (C) Quantification of the size. The major axis of the CENP-T shell-like and CENP-C structures were measured and plotted. Horizontal red lines indicate mean value. Statistical significance was tested using unpaired two-tailed Welch’s *t*-test. (D) Localization of CENP-N. DT40 cells expressing CENP-N-FLAG and CENP-C-GFP were immunostained with anti-FLAG and anti-ggCENP-T antibodies to visualize CENP-N and CENP-T (upper), or with anti-FLAG and anti-GFP antibodies to visualize CENP-N and CENP-C (lower). Representative ExM images are shown as in panel A. Scale bar, 2 µm in ExM. (E) Localization of CENP-A. cKO-CENP-C/GFP-CENP-C cells were treated with doxycycline for 48 h and subsequently immunostained with GFP-booster and anti-CENP-A antibodies. Representative ExM images are shown as in panel A. Scale bar, 2 µm in ExM. (F) Model of interphase kinetochore structure in DT40 cells.

Because the CENP-T staining profile that forms a spherical, shell-like structure may be a technical artifact, we tested a series of experimental conditions to rule out this possibility. First, we suspected that the CENP-T antibody’s characteristic recognition property could result in shell-like staining. To investigate this possibility, we examined the recognition site of the anti-CENP-T antibody via western blotting (Fig. S4A). The antibody recognized the entire CENP-T region, suggesting that the shell-like staining was not due to antibody specificity. Second, we considered the effect of tagging. Since we used mScarlet-HA-SNAP-CENP-T for CENP-T staining in DT40 cells, we examined whether endogenous CENP-T exhibited a similar structure in wild-type cells (Fig. S4B). A comparable spherical, shell-like structure was observed, suggesting that tagging was not responsible. Third, we suspected poor antibody penetration. To address this possibility, we immunostained cells expressing mScarlet-HA-SNAP-CENP-T using anti-CENP-T and anti-HA antibodies. Both antibodies revealed similar spherical, shell-like structures (Fig. S4C). Since the kinetochore core was accessible to the anti-GFP antibody, as demonstrated by CENP-C staining, it is unlikely that the shell-like structure resulted from limited antibody penetration. Finally, we applied an alternative ExM procedure: ultrastructure ExM (U-ExM), which has been reported to better preserve native structures (*39*). A similar spherical, shell-like structure was observed again under U-ExM in combination with deconvolution or super-resolution microscopy (Fig. S4D). Based on these analyses, we conclude that the CENP-T shell-like structure is genuine. Together, our results suggest that the CENP-T shell-like structure with a CENP-C core is a genuine feature of the kinetochore structure.

Next, we analyzed the localization of CENP-N, another CCAN component that binds to CENP-C via the CCAN-binding domain located in the N-terminal region (aa 166–324). We generated DT40 cells in which the endogenous CENP-N and CENP-C were replaced with CENP-N-FLAG and CENP-C-GFP, respectively. Then, we investigated the localization of CENP-T with CENP-N and CENP-C with CENP-N using ExM (Fig. 3D). CENP-N co-localized with CENP-T, exhibiting a shell-like profile (Fig. 3D, upper panel). In contrast, the C-terminal region of CENP-C was positioned internally and adjacent to CENP-N (Fig. 3D, lower panel). These observations are consistent with previous structural models, which suggest that the N-terminal region of CENP-C contacts CENP-L/N and that CENP-T directly contacts CENP-H/I/K/M and CENP-L/N (*21, 22*). Since the CENP-C C-terminus binds to the CENP-A nucleosome, we examined the localization of CENP-A next. We found that CENP-A co-localizes with CENP-C (Fig. 3E), suggesting that CENP-A resides at the CENP-C core. Additionally, we note that the kinetochore forms within a DAPI-sparse region surrounded by DAPI-dense chromatin (Figs. 3A, B, D, and E), as suggested by cryo-electron tomography analysis (*40*). Because CENP-A is localized at the core, centromeric chromatin appears to extrude from the surrounding heterochromatin and be incorporated into the kinetochore core.

Based on the ExM analyses, we summarized the localization profile of the CCAN proteins in Fig. 3F. We concluded that the interphase kinetochore in DT40 cells forms a spherical shell structure, in which the CENP-C core is surrounded by the CCAN complex. This finding is consistent with our MD simulation model.

### CENP-C is required for formation of the spherical shell-like structure in interphase DT40 cells

CENP-C is located at the core of the spherical shell structure of the CCAN, and our simulations suggest that it is crucial for aligning CCAN units (Figs 1 and 2). Therefore, we conducted an experiment to test how CENP-C contributes to the formation of the CCAN structure. To do so, we observed CENP-T localization in tet-inducible cKO-CENP-C DT40 cells using ExM. In the presence of CENP-C (no tet addition), we observed a spherical, shell-like CENP-T structure (Fig. 4, CENP-C +). The average size of the shell-like structure was 2.03 μm in ExM (Fig. 4, right). In contrast, the shell-like structure of CENP-T was abolished in the absence of CENP-C (tet addition), and CENP-T signals were aggregated at the center. The kinetochore size was significantly smaller than in cells expressing CENP-C (no tet addition) (Fig. 4, cKO-CENP-C). These results support the idea that CENP-C is required for forming the spherical, shell-like CCAN structure in DT40 cells.

**Figure 4.**
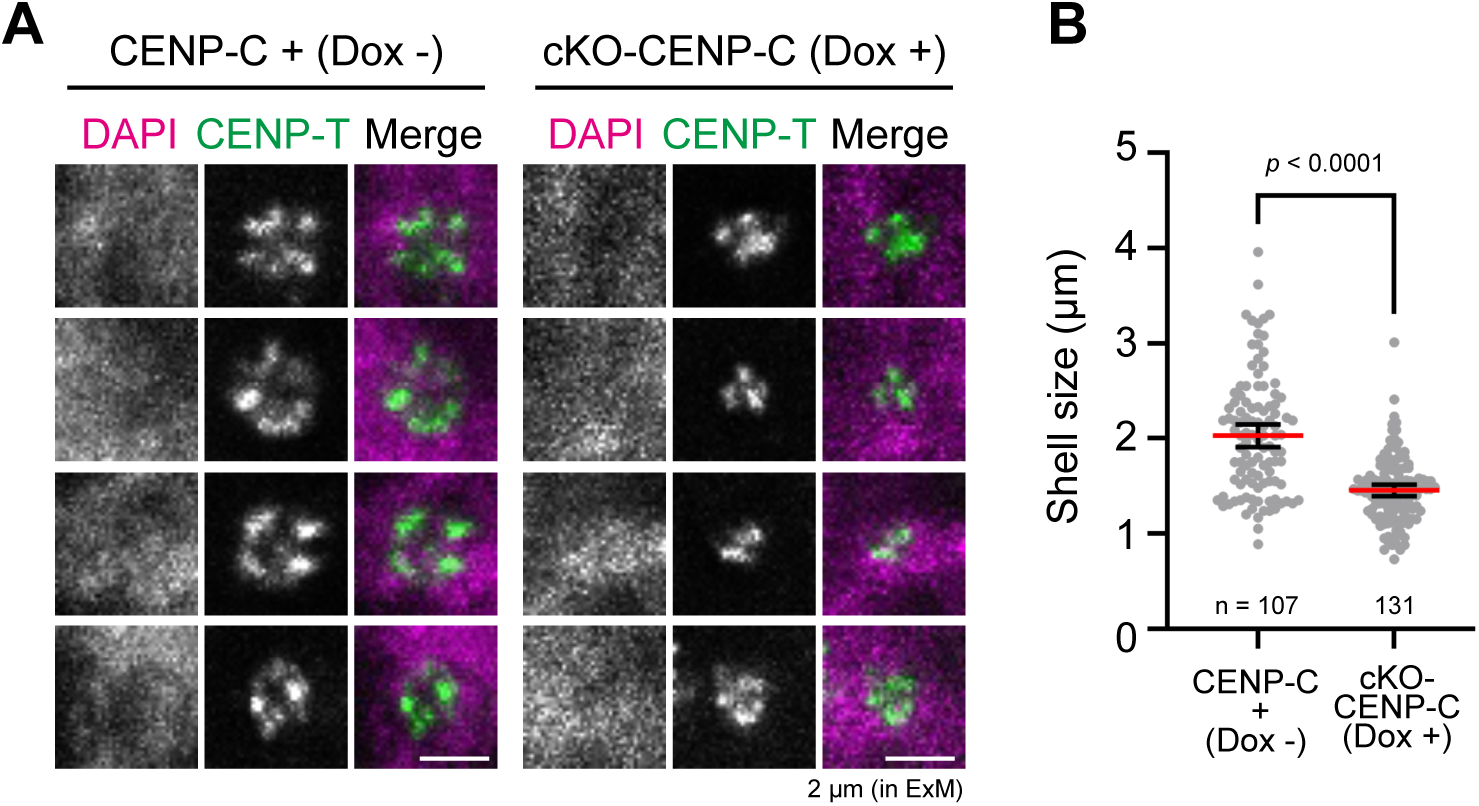
CENP-C is required for shell-like structure formation in DT40 cells. (A) CENP-T structure in the presence or absence of CENP-C. cKO-CENP-C was treated with (cKO-CENP-C) or without (CENP-C +) doxycycline for 48 h, and immunostained with anti-CENP-T antibody. Representative ExM images are shown. Scale bar, 2 µm in ExM. (B) Quantification of shell size. The major axis of the shell was measured and plotted. Horizontal lines indicate mean ± 95% confidence interval. Statistical significance was tested using unpaired two-tailed Welch’s *t*-test.

## Discussion

CCAN exists in tens of copies per kinetochore within cells. While various cryo-EM single particle analyses have provided structural details of a single CCAN unit, understanding how multiple CCAN units assemble within cells to form functional kinetochores remains one of the most central challenges in centromere/kinetochore biology. In this study, we addressed this issue by combining MD simulations with high-resolution imaging. Our data demonstrate that multiple CCAN units form a spherical shell-like structure in interphase kinetochores, in which the CENP-C core is surrounded by the CCAN subcomplexes.

This characteristic structure is not observed in the mitotic kinetochore, which instead adopts a disk-like plate structure, as seen under an electron microscope or an Expansion microscope (*38, 41*). These observations suggest that the shell-like structure forms only during interphase. However, a previous study reported that outer kinetochore proteins sometimes form a ring-like structure in mitotic kinetochores in *Xenopus* cells, though this structure is disrupted by microtubule depolymerization (*42*). The authors proposed that the ring-like structure may transform into a disk-like structure when microtubules attach to the kinetochores, properly. Since various outer kinetochore proteins bind to CENP-T (*6, 29, 43, 44*), the spherical, shell-like CCAN structure observed by us may form the basis for outer kinetochore assembly. Considering the function of the kinetochore, efficient microtubule capture during the M phase requires the formation of a highly organized disk-like structure. To prepare for this transition, the kinetochore may adopt a pre-organized, bundle-like structure of CCAN units during interphase. Consistent with our observations, a similar shell-like structure has been also observed in *Xenopus* cells (*45*).

Our results demonstrate that CENP-C plays a pivotal role in establishing the shell-like structure; this structure becomes disorganized in the absence of CENP-C. Consistent with these observations, Mini-CENP-C localizes to kinetochores during interphase but is almost absent from the mitotic kinetochore, whereas the mitotic kinetochore remains functional (*31*). These finding suggest that CENP-C contributes only minimally to mitotic kinetochore architecture. However, mutations that impair CENP-C oligomerization result in mitotic abnormalities, indicating that the CENP-C-mediated shell-like structure of the CCAN may be essential for the functional kinetochore formation upon entry into M-phase. Alternatively, the shell-like structure could be involved in interphase-specific events such as DNA replication and transcription in the centromere region. Further experiments are required to clarify the significance of this interphase-specific structure. Interestingly, a similar structure has also been observed in *Xenopus* kinetochores (*45*). These observations suggest that this shell-like organization represents a conserved architectural feature of interphase kinetochores. Understanding the significance of this structure will provide important insights into the functional and structural principles of the kinetochore.

In cryo-EM structures of the human CCAN-DNA complex, CENP-L/N forms a tunnel that grips the DNA strand extending from the CENP-A nucleosome core (*21, 22*). CENP-C contains the conserved CENP-L/N binding motif, which is followed by a basic stretch (Fig. 1A). Our MD simulations of the 2(CCAN-DNA)-Mini-CENP-C dimer complex suggest that the basic stretch of CENP-C transiently interacts with the DNA as it enters the CENP-L/N basic tunnel (Fig. 1H). Such interactions, which are mediated by electrostatic contacts between the basic residues of CENP-C and the DNA phosphate backbone, were not observed in the cryo-EM structures. This is likely due to their transient and dynamic nature. Nevertheless, these interactions are likely to contribute significantly to the formation of higher-order CCAN assemblies (Fig. 2). Simultaneous interactions of CENP-C with both CENP-L/N and DNA may anchor the CCAN to chromosomal DNA and constrain the mobility of the CCAN-DNA complex within chromatin. Consistent with this idea, simulations of the 8(CCAN-DNA)-4Mini-CENP-C dimer complex revealed that all DNA fragments bound to individual CCAN units were oriented toward the CENP-C based center of the spherical, shell-like structure. In contrast, DNA fragments in a complex containing the Mini-CENP-C^6E^ mutant exhibited random orientations (Fig. 2A).

These findings suggest that the formation of the CCAN shell-like structure require DNA binding by CENP-C through its conserved basic stretch near the CENP-L/N binding site (Fig. 2). We propose that this DNA interaction stabilizes the association between CENP-C and the CENP-L/N subcomplex and thereby promotes ordered CCAN assembly. Mutations within this region (the 3E mutation) cause mitotic defects, suggesting that disruption of CENP-C – DNA interactions compromises CCAN organization in cells. Considering that CENP-A is located within the shell-like structure in a DAPI-sparse region, appropriate contacts between CENP-C and DNA are likely essential for forming and maintaining the higher-order CCAN assembly.

Here, we demonstrated a high-resolution structure using ExM techniques. Recently, other approaches, such as cryo-electron tomography, have become available for analyzing high-resolution structures of supramolecular complexes in cells (*40, 46–49*). However, the limited number of kinetochore copies does not allow for an efficient averaging process to obtain its high-resolution structure using cryo-electron tomography (*40*). In such cases, combining MD simulations with high-resolution fluorescence imaging provides a powerful strategy for investigating mesoscale cellular structures between individual molecules (nanoscale) and whole cells (microscale). Using this integrated approach, we revealed the structure of the kinetochore. These findings provide important insights into the spatial organization of the functional kinetochore within cells.

## Methods

### Initial model generation for MD simulations

Structural models of the CCAN unit and the dimeric CENP-C cupin-containing region (CENP-C cupin) were generated using the AlphaFold3 (AF3) prediction server (DeepMind/Isomorphic Labs; https://alphafoldserver.com/). The CCAN unit components used for each prediction are summarized in Supplementary Tables S1-S3. For each prediction, five models were generated and ranked based on the predicted local distance difference test (pLDDT) score. Model confidence was further evaluated using the predicted aligned error (PAE) and, for multimeric predictions, the interface predicted template modeling (ipTM) score. The highest-confidence model was selected for downstream model building of the CCAN unit–Mini-CENP-C complex. The unstructured regions with low confidence scores were eliminated from the initial model (Table S2). Model visualization, confidence assessment, and subsequent model manipulation were performed in UCSF ChimeraX (*50*).

To obtain full-length dimeric CCAN-Mini-CENP-C models, two CCAN unit models and one CENP-C cupin dimer were positioned approximately 180 Å apart. This distance was chosen to ensure sufficient spatial separation within a defined modeling volume. Missing regions corresponding to the Mini-CENP-C intrinsically disordered segments linking the CCAN-binding and cupin-dimer regions were generated using MODELLER through the ChimeraX interface (*51*). In total, six CCAN_mini-CENP-C dimer models with distinct CCAN orientations were generated.

Complex models containing eight CCAN units were assembled using AF3-generated CCAN core and dimeric CENP-C cupin models. Four CENP-C cupin dimers were arranged at the center of a 50 × 50 × 50 nm cubic volume based on the oligomeric interface observed in the CENP-C cupin crystal structure (PDB: 7X85). Eight CCAN unit models were then placed around the central cupin assembly within the cubic space. Missing mini-CENP-C IDRs in these octameric assemblies were generated using MODELLER via ChimeraX (*51*).

### MD Simulations

The CG simulations were performed using the GROMACS (version 2022) (*52*) and the MARTINI model (version 2.2) (*33, 34*). The initial structure of the protein was predicted using AF3 (*32*). In the CG model, the tertiary structures of proteins excluding the Mini-CENP-C IDR regions were maintained using an elastic network model. The proteins and DNAs were solvated in a 70 nm × 70 nm × 70 nm box with CG waters and 0.15 M NaCl ions. The initial configurations were built by the Martini Maker module in CHARMM-GUI server (*53*). Each system was first energy minimized and equilibrated using the Berendsen thermostat and barostat (*54*) followed by 1 μs production runs with a 20 fs time step. System temperature and pressure during the production phase were maintained at 303.15 K and 1 atm with the velocity rescaling thermostat and the semi-isotropic Parrinello–Rahman barostat (*55*), respectively.

The all-atom simulations were performed using GENESIS (*56*) and the CHARMM36 force-field (*57*). The initial configurations were built by the Solution Builder module in CHARMM-GUI server (*53*). Energy minimization was performed for 1000 steps by the steepest descent algorithm and then by the conjugate gradient algorithm. Then a 250 ps NVT simulation was performed at 303.15 K for solvent equilibration, followed by a 1.6 ns NPT equilibration to 1 atm using the Langevin thermostat/barostat (*58*). MD simulations were performed for 100 ns with a time-step of 2 fs and Langevin thermostat/barostat. Long-range electrostatic interactions were treated by the particle-mesh Ewald method (*59, 60*). The short-range electrostatic and van der Waals interactions both used a cutoff of 12 Å. All bonds were constrained by the SHAKE/RATTLE algorithm (*61, 62*).

The relative orientation of two CCANs was defined as the direction cosine of the two DNA directional vectors within each CCAN (with the direction from CCAN to cupin domain of Mini-CENP-C defined as positive). K232 in CENP-L and K261 in CENP-I were stably bound to DNA at the most distant positions. In the absence of DNA, the relative orientation was defined as the direction cosine of the direction vectors of these residues. The simulation results were visualized and analyzed using PyMOL (http://www.pymol.org/pymol).

### DNA Construction

mScarlet-HA-SNAP-CENP-T DNA sequence was amplified by PCR and cloned into a pBluescript KS(+) backbone and flanked by ∼1 kb homology arms surrounding the CENP-T start codon, using In-Fusion system (Clontech, Cat. No. 638948) according to manufacturer’s protocol.

To generate GFP-Mini-CENP-C^3E^ mutant, GFP-Mini-CENP-C plasmid (*31*) was inversely amplified using primers, which introduced 3E mutation (K216E/R223E/R226E), and the PCR fragments were circularized by In-Fusion system.

To express GST-fused chicken CENP-T fragments in *Eshcerichia coli*, DNA sequences corresponding to aa 1-150, 151-250, 251-500, and 501-639 were amplified by PCR and then inserted into the BamHI- and XhoI-digested pGEX-6P-1 vector (Cytiva, Cat. No. 28954648) using In-Fusion system.

All cloned DNA sequences were confirmed by the Sanger sequence.

### Chicken DT40 cells

A chicken DT40 cell line CL18 was used as the wild-type (WT) cell (*63*). DT40 cells were cultured at 38.5°C in DMEM medium (Nacalai Tesque) supplemented with 10% fetal bovine serum (FBS; Sigma), 1% chicken serum (Thermo Fisher Scientific), and Penicillin-Streptomycin (Thermo Fisher Scientific, Cat. No. 15140122) (DT40 culture medium).

The chicken CENP-C conditional knockout (cKO-CENP-C) DT40 cell line was described before (*37*). The cKO-CENP-C cells expressing GFP-Mini-CENP-C^3E^ were generated by random integration of the linear plasmid construct. The cKO-CENP-C cell line expressing GFP-Mini-CENP-C (*29*) or Mini-CENP-C^3E^ were used. To conditionally knockout CENP-C, the cells were cultured in DT40 culture medium containing 2 μg/ml tetracycline (Tet, Sigma, Cat. No. 87128) for 48-72 h.

DT40 cells in which endogenous CENP-T or CENP-N was replaced with mScarlet-HA-SNAP–CENP-T or CENP-N-FLAG, respectively were generated using CRISPR/Cas9 system. Guide RNA and homology arm sequences for them have been previously reported (*31*).

### Immunostaining

DT40 cells were attached to coverslips using Cytospin3 at 800 rpm for 5 min. Cells were fixed with 4% paraformaldehyde (PFA; Electron Microscopy Sciences, Cat. No. 15710) in 250 mM Hepes-NaOH (pH 7.4) for 10 min at room temperature (RT; 25–28 °C), except for CENP-A staining. For CENP-A, cells were fixed with 4% PFA in phosphate-buffered saline (PBS) for 1 min at RT, followed by methanol fixation at –30 °C for 20 min. After blocking with blocking buffer (10% Blocking One [Nacalai Tesque, Cat. No. 03953-95], 0.5% Triton X-100 in PBS) for 30 min, cells were incubated with primary antibodies for 2 h at RT: anti-ggCENP-T (*11*) (1:1,000), anti-ggCENP-A (*11*) (1:500), anti-GFP (MBL, Cat. No. 598; 1:1,000), and anti-FLAG M2 (Sigma, Cat. No. F3165; 1:1,000), anti-HA (Roche, Cat. No. 12158167001; 3F10, 1:1,000), anti-α-tubulin (DM1A, Sigma, Cat. No. T6199; 1:1000), GFP-booster-Alexa488 (chromotek, Cat No. gb2AF488; 1:200). After three washes, cells were incubated with appropriate secondary antibodies together with 100 ng/mL 4′,6-diamidino-2-phenylindole (DAPI; Sigma-Aldrich, Cat. No D9542): anti-rabbit IgG–Alexa488 (Thermo Fisher Scientific, Cat. No. A-11034, 1:250), anti-rabbit IgG–Alexa568 (Thermo Fisher Scientific, Cat. No. A-11036, 1:250), anti-mouse IgG–Alexa488 (Thermo Fisher Scientific, Cat. No. A-11029, 1:250), and anti-mouse IgG–Alexa568 (Thermo Fisher Scientific, Cat. No. A-11031, 1:250), anti-rat IgG-Alexa568 (Thermo Fisher Scientific, Cat. No. A-11077, 1:200). Following washing, cells were directly subjected to expansion microscopy.

### Cell counting

To count DT40 cell numbers, the culture was mixed with the same volume of 0.4% (w/v) Trypan Blue Solution (FUJIFILM, Cat. No. 594-31481) and the cell numbers were counted by Countess II (Thermo Fisher Scientific).

### Quantification of mitotic index

The cKO-CENP-C cells expressing GFP-Mini-CENP-C or GFP-Mini-CENP-C^3E^ were cultured in DT40 culture medium in the presence or absence of 2 μg/ml tetracycline for 72 h at 38.5°C, and then immunostained with an anti-α-tubulin antibody as described above. The images were acquired every 0.2 μm intervals of z-slice using a Fusion BT sCMOS camera (HAMAMATSU photonics) mounted on a Nikon Eclipse Ti inverted microscope with an objective lens (Nikon; Plan Apo lambda 100x/1.45 NA) with a spinning disk confocal scanner unit (CSU-W1; Yokogawa) controlled with NIS-elements (Nikon; version 5.42.01). The images in the figures are the maximum intensity projection of the Z-stack generated with Fiji (*64*). The mitotic index (mitotic cell numbers / total cell numbers) was manually counted. Statistical analysis of the measurements was performed using GraphPad Prism software (version 10.4.1 or later).

### Expansion microscopy (ExM)

We used 12× ExM as reported by Norman et al. (*38*), unless otherwise noted, with slight modifications. Briefly, immunostained cells were post-fixed with 2% glutaraldehyde (GA; EM grade, Electron Microscopy Sciences, Cat. No. 16220) in PBS overnight at RT. After repeating GA fixation for 15 min, cells were embedded in hydrogel containing N, N-dimethylacrylamide (Sigma-Aldrich, Cat. No. 274135) and sodium acrylate (SA; AK Scientific, Cat. No. R624-25g), digested with 30 units of proteinase K (Sigma-Aldrich, Cat. No. P4850) in digestion buffer (0.5 M SDS, 0.5% Triton X-100 in TAE) together with 100 ng/ml DAPI at 37 °C for 3 h, and expanded in Milli-Q water. The expansion factor measured by expanded gel size was ∼10. The expanded gel was trimmed to an appropriate size and mounted on a poly-L-lysine–coated 35 mm glass-bottom dish (Matsunami, Cat. No. D11141H). Images were acquired using an AxioObserver.Z1 system (Carl Zeiss) equipped with a water-immersion 63× C-Apochromat objective lens (NA = 1.2) and a Prime BSI sCMOS camera (Teledyne). Fluorescence excitation was performed with an LED light source (X-cite Xylis2), and emission was separated using appropriate dichroic mirrors and filters: FT 395, BP 445/50 for blue; FT 495, BP 525/50 for green; FT 570, BP 605/70 for red fluorescence. The microscope was controlled with ZEN blue 3.6 software (version 3.6.5). Raw data were deconvolved using ZEN, and when sample drift was prominent, drift correction was applied prior to deconvolution using 3D-Aligner (*65*).

To confirm the shell-like structure of CENP-T, we used U-ExM (*39*) in combination with super-resolution microscopy. Immunostained cells were post-fixed with fixation buffer (1.4% PFA, 2% acrylamide [Sigma, Cat. No. A4058]) overnight at RT. Cells were then embedded in monomer solution (10% acrylamide, 0.1% methylenebisacrylamide [Sigma, Cat. No. M1533], 19% SA in PBS), denatured in denaturation buffer (50 mM Tris-HCl [pH 9.0], 200 mM NaCl, 200 mM SDS) together with 100 ng/mL DAPI at 95 °C for 3 h, and subsequently expanded in Milli-Q water. The expanded gel was mounted on a poly-L-lysine–coated 35 mm glass-bottom dish. Imaging was performed using either an LSM880 with Airyscan (Carl Zeiss) equipped with a 63× C-Apochromat objective lens or a Deltavision Elite system (GE Healthcare) equipped with a silicone-immersion 60× UPlanSApo objective lens (NA = 1.3; Olympus). Both microscopes were operated with their default control software (ZEN black 2.3 SP1 FP2, version 14.0.0.0; SoftWoRx, version 7.0.0). Airyscan processing was performed using default settings. Enhanced ratio deconvolution was applied in SoftWoRx using default settings with a custom optical transfer function of the 60× UPlanSApo objective lens.

Image contrast was adjusted in Fiji/ImageJ without altering gamma values for optimal presentation.

### Quantification of shell size

The size of the CENP-T shell was quantified using Fiji/ImageJ. The major axis of each shell was manually selected and measured. Statistical analysis of the measurements was performed using GraphPad Prism software (version 10.4.1 or later).

### Protein purification and western blotting

GST-fused CENP-T fragments were expressed in BL21 *E. Coli* cells. Cells harboring the plasmids were cultured in Luria Broth medium at 37°C until OD600 of ∼0.6. Protein expression was induced by adding 1 mM isopropyl-β-D-thiogalactopyranoside (IPTG), and the cultures were incubated for an additional 3 h at 25 °C.

Cells were harvested, washed with PBS, and resuspended in PBS containing 0.5% Trition-X-100 and cOmplete protease inhibitor cocktail (Roche, Cat. No. 4693116001). After sonication, the extract was clarified by centrifugation at 12,000 x g for 10 min at 4°C. The supernatant was incubated with glutathione sepharose 4B bead for 1 h at 4°C. After three washes with PBS containing 0.5% Trition-X-100, the purified GST-CENP-T fragments were analyzed by SDS-PAGE and CBB staining using BSA as a standard. Five hundred ng or 50 ng of GST-CENP-T fragments were used for CBB staining and western blotting, respectively. Proteins were transferred to a PVDF membrane, probed with anti-ggCENP-T (1:2000) and anti-GST (Nacalai Tesque, Cat. No. 04435-84, 1:5000) antibodies, and detected by chemiluminescence (ImmunoStar Zeta, Fujifilm Wako, Cat. No. 297-72403).

## Supplemental Information

The article contains four Supplemental Figures, three Supplemental Tables and seven Supplemental Videos.

## Acknowledgments

The authors are very grateful to members of the Fukagawa Lab for the fruitful discussion. We also thank R. Fukuoka and Y. Kubota for technical assistance. YS, MA, and YH equally contributed to this work. The order of the authors on the list corresponds to the order in which they created the figures, enabling each author to change the order and list their name first on their CV. The numerical computations were performed with the supercomputer systems Fugaku (hp230252, hp250019) and HOKUSAI at RIKEN, Wisteria/BDEC-01 at the University of Tokyo. This work was supported by CREST of JST (JPMJCR21E6), JSPS KAKENHI Grant Numbers 24H02280, 24H02281, and 25H00975 to TF, JSPS KAKENHI Grant Numbers 23K05715, 24K01980, and 25K09559 to YS, JSPS KAKENHI Grant Numbers 25K09518, and 24H02286 to YH, JSPS KAKENHI Grant Number 24K09340 to MA.

## Author contributions

Conceptualization: Y.S., M.A., Y.H., T.F.; Investigation: Y.S., M.A., T.H., A.S., T.H., M.T., T.F.; Formal analysis: Y.S., M.A., Y.H., T.F; Resources: Y.S., M.A., Y.H., T.F.; Data curation: Y.S., M.A., Y.H., T.F.; Writing - original draft: Y.S., M.A., Y.H., T.F.; Writing - review & editing: T.H., Y.M., M.A., T.F.; Supervision: T.F.; Project administration: T.F.; Funding acquisition: Y.S., M.A., Y.H., T.F.

## Supplementary Figure legends

**Supplementary Fig S1.**
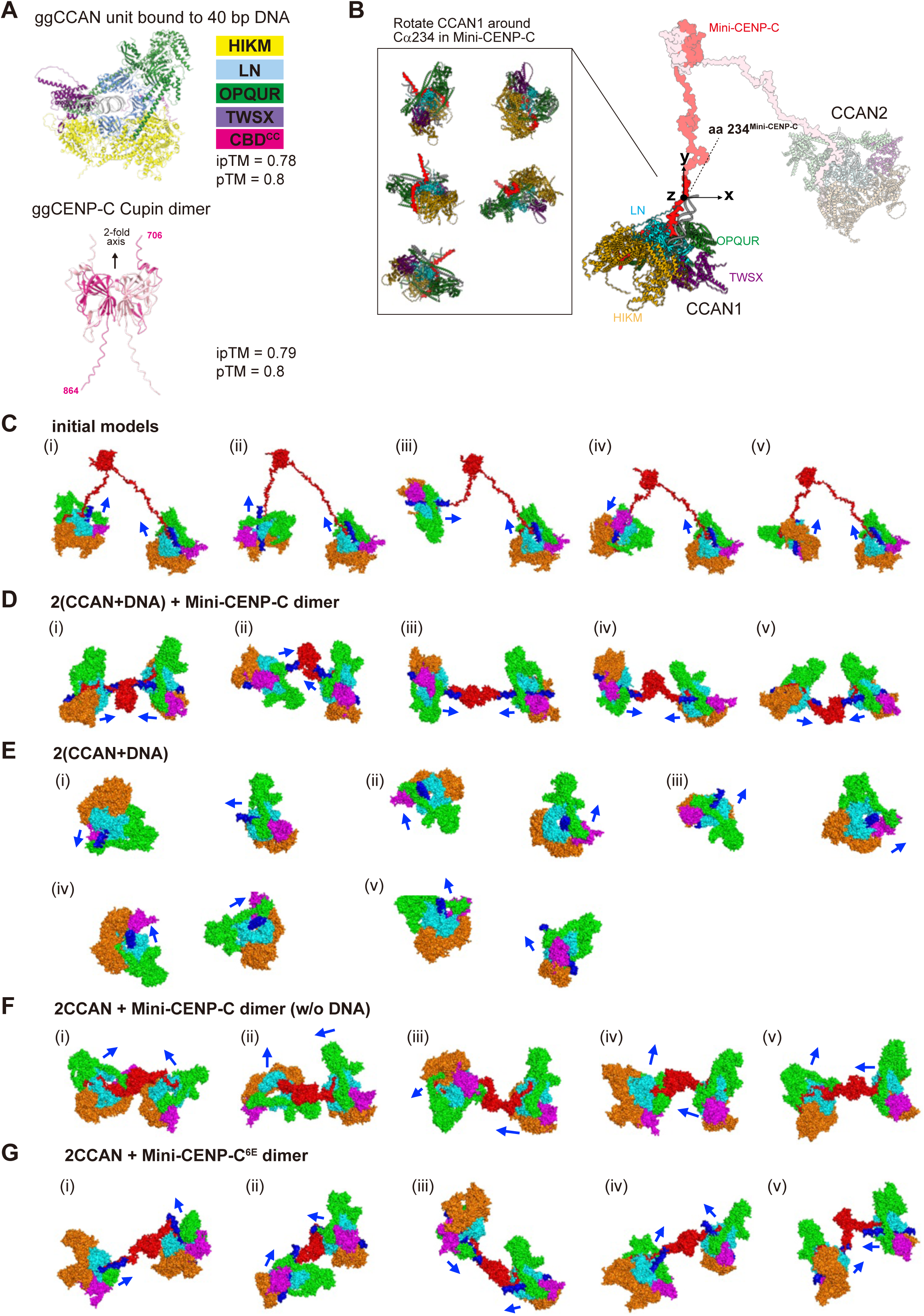
MD simulation of two CCAN complexes with DNA linked by a Mini-CENP-C dimer. (A) AlphaFold3-predicted structural models used for initial model generation for simulations. Top, The model of the chicken CCAN unit. The CCAN subcomplexes, CENP-H/I/K/M, -L/N, -O/P/Q/U/R, and -T/W/S/X, are shown in the indicated color code. The model contains a 40 base pair double stranded DNA fragment (light gray) and the N-terminal half of CENP-C CCAN Binding Domain (CBD; magenta). Bottom, The model of the dimeric cupin domain of chicken CENP-C (amino acids 706-864). Regions predicted with low confidence were excluded from the models (see Methods). (B) Generation of initial models for MD simulations. Two DNA-bound CCAN units (CCAN1 and CCAN2) were tethered by a Mini-CENP-C dimer. To generate distinct initial relative orientations between two CCAN units, CCAN 1 was systematically rotated around the main-chain Cα atom of residue 234 in Mini-CENP-C as illustrated. Five additional initial models with different orientations of the CCAN1 unit (left panel) were generated. (C) Five initial models of two DNA-bound CCAN units linked by a Mini-CENP-C dimer (i-v). The models were generated as described in (B) and subjected to CG MD simulation shown in (D). (D) Structures of the full complexes [2(CCAN+DNA) + Mini-CENP-C dimer] at 500 ns in CG MD simulations initiated from different initial models shown in Fig. S1C. (E) Structures of two DNA-bound CCAN units simulated in the absence of Mini-CENP-C at 500 ns. (F) Structures of two CCAN units (without DNA) linked by a Mini-CENP-C dimer at 500 ns. (G) Structures of two DNA-bound CCAN units inked by Mini-CENP-C basic patch mutant, Mini-CENP-C^6E^ at 500 ns. For each panel, (i–v) correspond to independent simulations starting from the respective initial states shown in panel (C). Color scheme as in Fig. 1B.

**Supplementary Fig S2.**
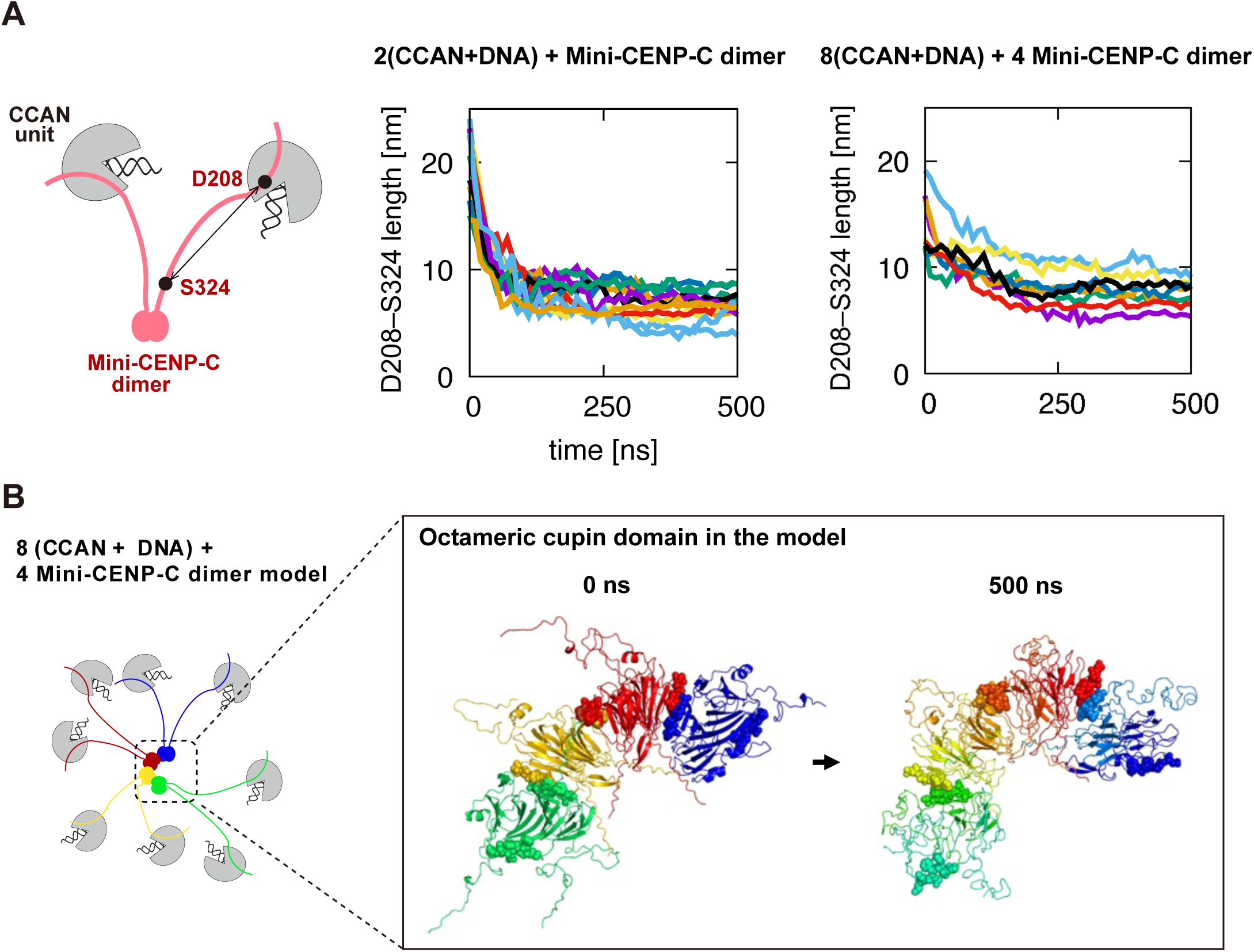
Structural convergence of the CCAN-DNA-Mini-CENP-C complex during MD simulation. (A) Temporal changes of the length of the intrinsically disordered region (IDR) of Mini-CENP-C during CG MD simulations of CCAN-DNA-Mini-CENP-C complexes. Left, schematic presentation showing the definition of IDR length, measured as the distance between Mini-CENP-C residues D208 and S324. Middle, simulation of two DNA-bound CCAN units tethered by dimeric Mini-CENP-C. The temporal changes of 12 IDR lengths (two per dimeric Mini-CENP-C unit) observed in six independent simulations. Right, temporal changes of IDR lengths in octameric Mini-CENP-C (four dimeric Mini-CENP-C). (B) Schematic representation of the initial model of eight DNA-bound CCAN units tethered by octameric Mini-CENP-C (left). The Mini-CENP-C octamer model was generated based on the dimer–dimer interface observed in the crystal structure of the chicken CENP-C cupin domain (PDB ID: 7X85). Right, close-up views of the octameric cupin domain in the initial configuration (0 ns) and after MD simulation (500 ns).

**Supplementary Fig S3.**
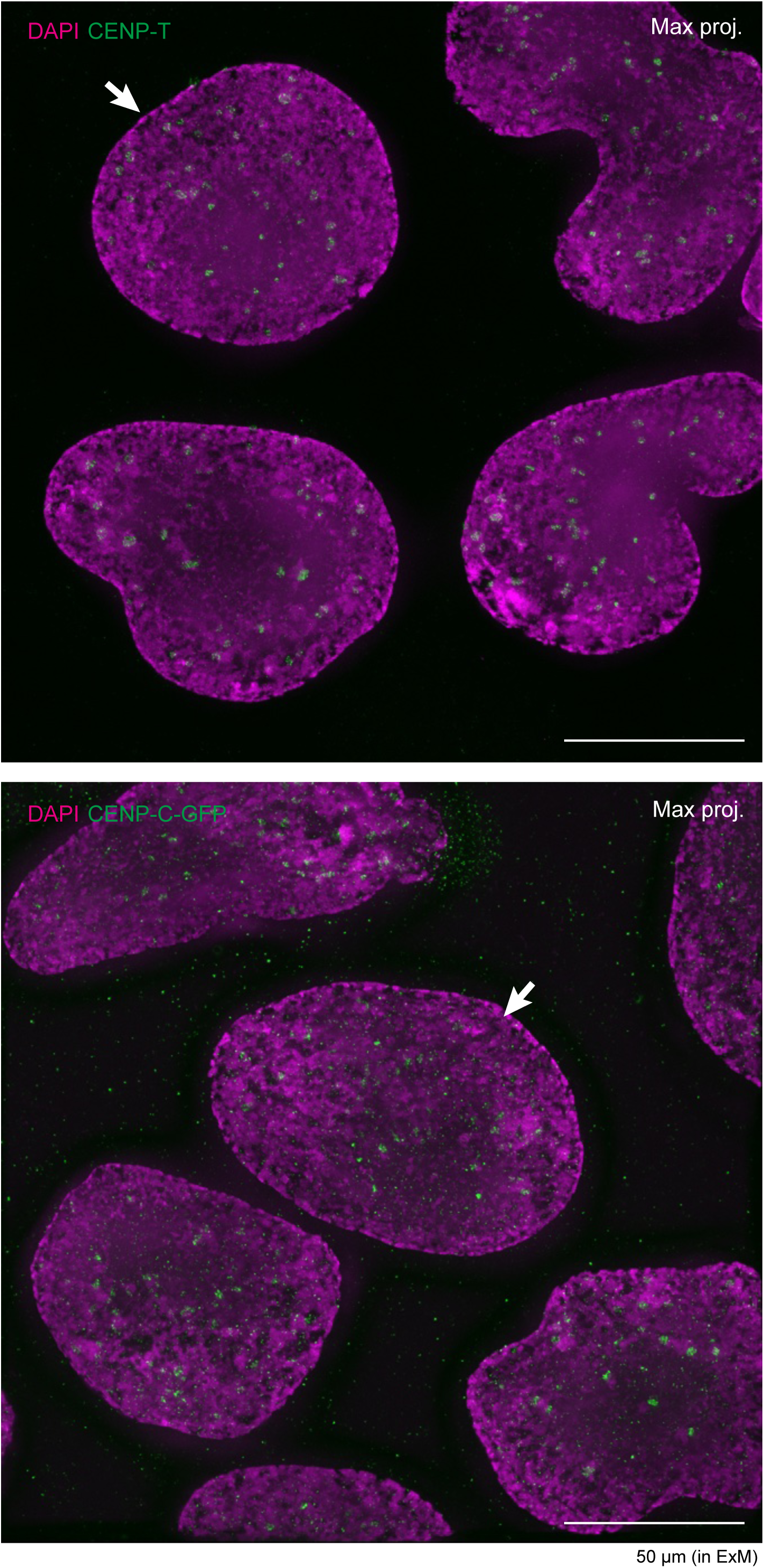
Wide-field view of ExM for CENP-T and CENP-C. Representative wide-field ExM images of CENP-T (upper) and CENP-C (lower) are shown. Cells indicated by white arrows correspond to those shown in Figure 3.

**Supplementary Fig S4.**
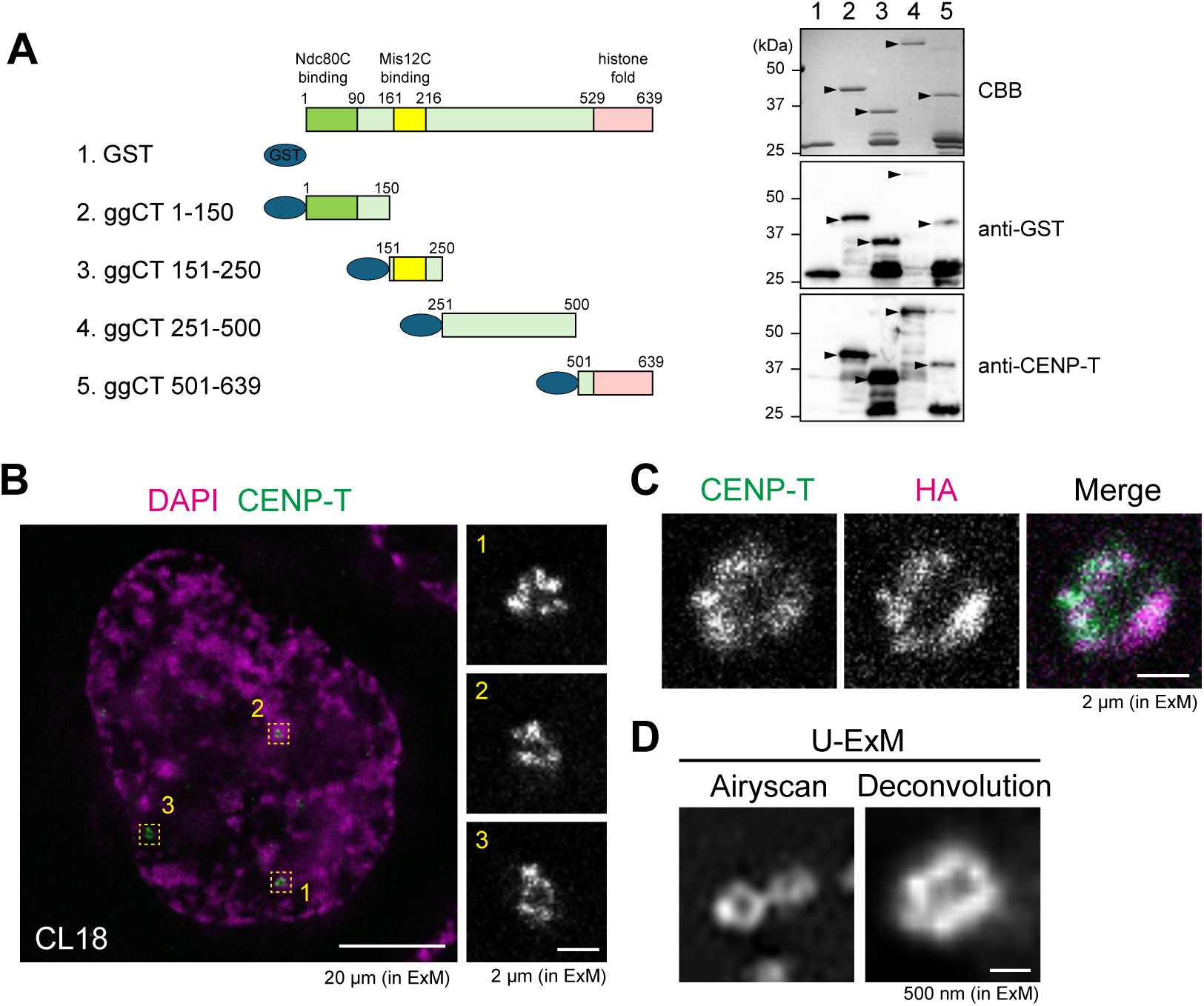
Shell-like CNEP-T structure was observed independently of experimental conditions. (A) Recognition site of anti-ggCENP-T antibody. GST-fused CENP-T fragments (left panel) were expressed and purified (right panel, CBB). The fragments were detected using either anti-GST or anti-CENP-T antibodies. Arrowheads indicate the full-length band. (B) Shell-like structure of endogenous CENP-T. Endogenous CENP-T was immunostained with anti-CENP-T antibody and observed by ExM. Deconvolved images are shown. Scale bars, 20 µm (left) and 2 µm (right) in ExM. (C) Co-staining of CENP-T with different antibodies. Cells expressing mScalret-HA-SNAP-CENP-T were immunostained with anti-CENP-T and anti-HA antibodies and observed by ExM. Deconvolved images are shown. Scale bar, 2 µm in ExM. (D) Observation using different ExM methods. Endogenous CENP-T was immunostained with anti-CENP-T antibody and observed by U-ExM in combination with Airyscan or deconvolution.

**Supplementary Table 1.**
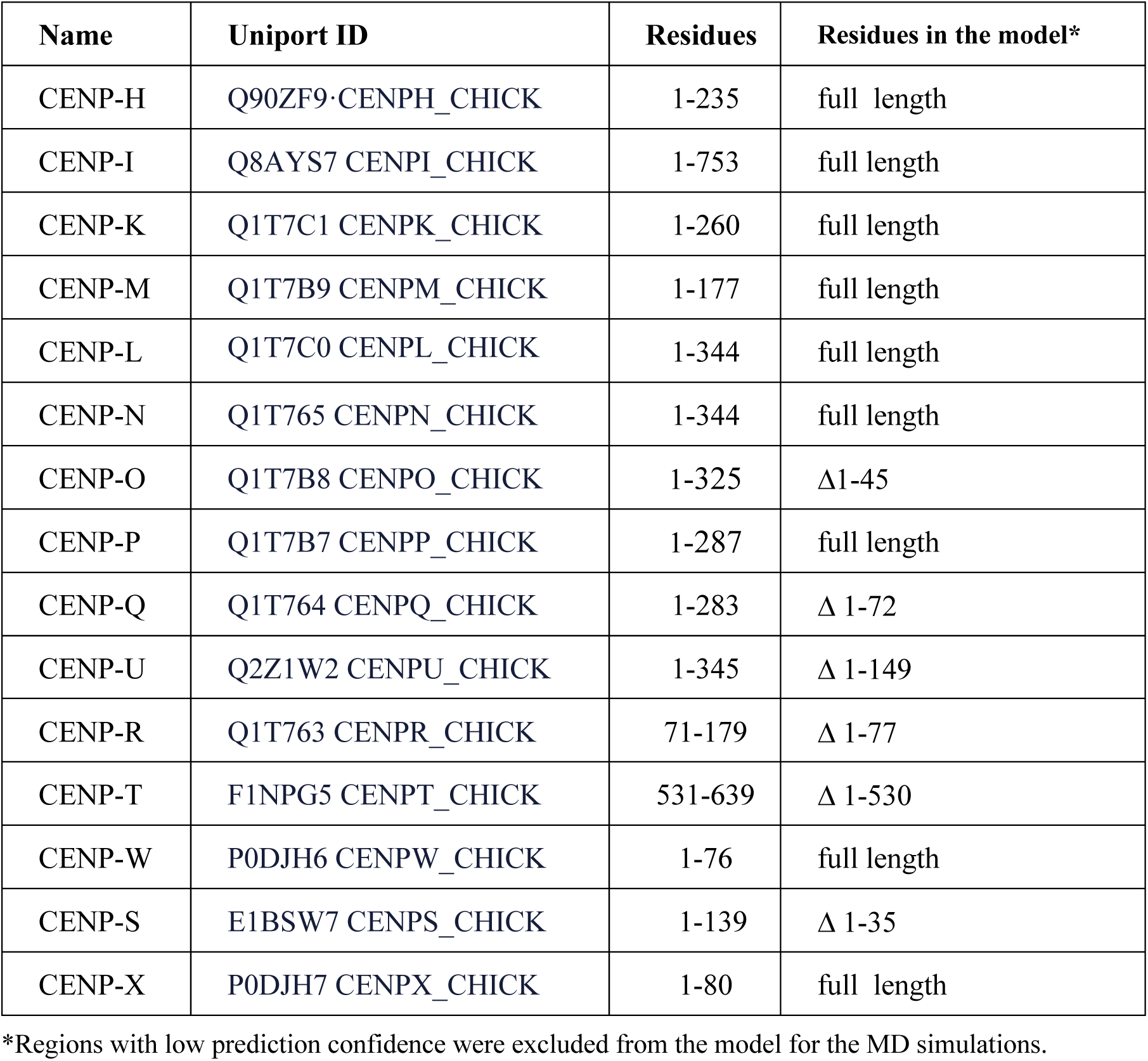
CCAN core components applied to AF3 prediction.

**Supplementary Table 2.**
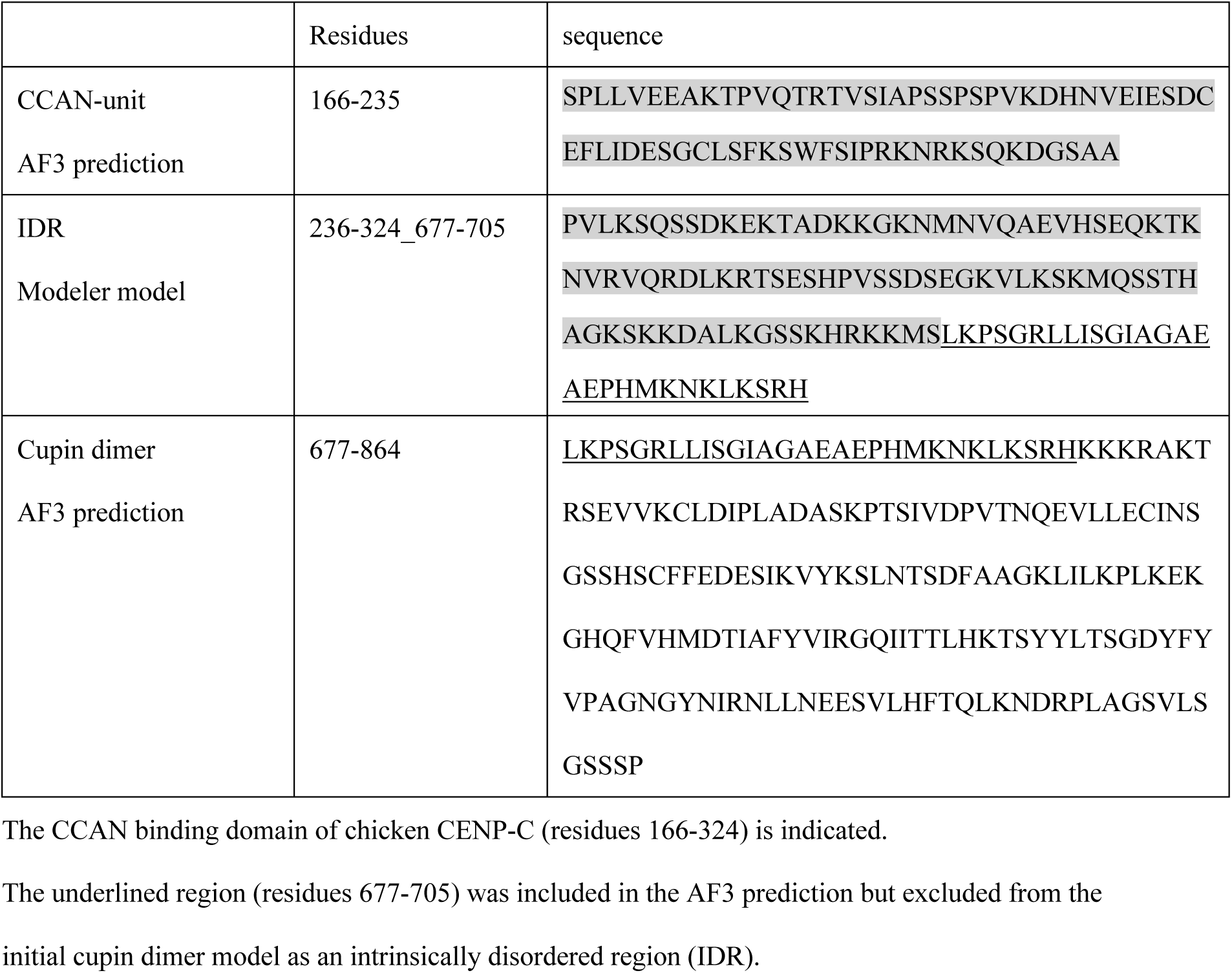
Mini-CENP-C sequence applied to initial model generation.

**Supplementary Table 3.**
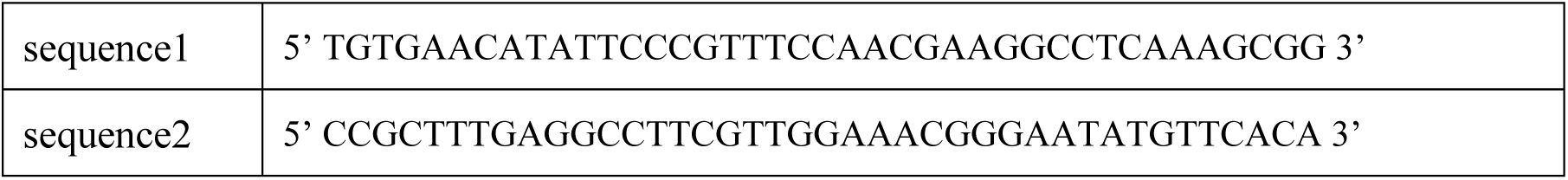
DNA sequences used in AF3 prediction.

**Supplementary Movie S1 MD simulation of two CCAN units with DNA linked by a Mini-CENP-C dimer**.

Corresponding to the snapshot in Fig. 1(B), with the same colour scheme.

**Supplementary Movie S2 MD simulation of two CCAN units with DNA in the absence of Mini-CENP-C dimer.**

Corresponding to the snapshot in Fig. 1(D), with the same colour scheme.

**Supplementary Movie S3 MD simulation of two CCAN units lacking DNA but linked by a Mini-CENP-C dimer.**

Corresponding to the snapshot in Fig. 1(F), with the same colour scheme.

**Supplementary Movie S4 MD simulation of two CCAN units with DNA linked by a Mini-CENP-C basic patch mutant dimer.**

Corresponding to the snapshot in Fig. 1(K), with the same colour scheme.

**Supplementary Movie S5 MD simulation of eight CCAN units with DNA in the absence of Mini-CENP-C**.

Corresponding to the snapshot in Fig. 2(A) upper panel, with the same colour scheme.

**Supplementary Movie S6 MD simulation of eight CCAN units with DNA linked by four Mini-CENP-C dimers**.

Corresponding to the snapshot in Fig. 2(A) middle panel, with the same colour scheme.

**Supplementary Movie S7 MD simulation eight CCAN units with DNA linked by four Mini-CENP-C basic patch mutant dimers**.

Corresponding to the snapshot in Fig. 2(A) bottom panel, with the same colour scheme.

